# Drivers of phylogenetic structure in Amazonian freshwater fish assemblages

**DOI:** 10.1101/2021.07.29.454320

**Authors:** Laís Salgueiro, Fernanda Cassemiro, James S. Albert, Renata G. Frederico, Max Hidalgo, Bernard Hugueny, Céline Jézéquel, Hernan Ortega, Pablo A. Tedesco, Gislene Torrente-Vilara, Jansen Zuanon, Thierry Oberdorff, Murilo S. Dias

## Abstract

**Aim:** Community phylogenetics provides important information about the evolutionary and ecological factors help structure regional species assemblages. Here, we analyze phylogenetic diversity (phylodiversity) patterns among fish species in 97 sub-drainages of the Amazon basin, to evaluate the roles of historical and contemporary processes in generating and maintaining the exceptional richness and endemism of Amazonian fish species assemblages.

**Location:** Amazon River basin

**Taxon:** Freshwater fishes

**Methods:** Using a large comprehensive database of freshwater fish species distributions, and a well-sampled molecular phylogeny of ray-finned (actinopterygian) fishes, we develop of multivariate statistical model to correlate estimated historical and contemporary environmental parameters with sub-drainage phylodiversity patterns. The model employs three phylogenetic metrics: i.e.: phylogenetic diversity (PD) *sensu stricto*, mean pairwise phylogenetic distance (MPD) between species capturing phylodiversity variation at older evolutionary timescales), and mean nearest taxon distance (MNTD) capturing variation in phylodiversity at younger evolutionary timescales.

**Results:** The model recovered significant effects of elevation gradients, contemporary climate, habitat fragmentation, water types, and past marine incursions on assemblage phylodiversity patterns. The model also found significantly negative relationships among the three phylogenetic metrics, and between these metrics and distance to mouth of the Amazon, representing a West-East longitudinal gradient.

**Main conclusions:** Our study revealed a highly non-random spatial and environmental distribution of our three phylogenetic diversity metrics across the 97 sub-drainages of the Amazon basin. Beyond significant regional effects of several environmental and historical drivers, we also found a significant West-East gradient of increasing phylogenetic diversity and phylogenetic relatedness, both patterns suggesting deeper evolutionary divergences among taxa located to the east, and more diverse, more recent radiations in the western sub-drainages. We conclude that western Amazonia can be seen as an evolutionary “cradle” of biodiversity for freshwater fishes in the Amazon basin as a whole.

**Significance Statement:** This manuscript reveals spatial patterns of freshwater fish phylogenetic diversity and relatedness and explains its major contemporary and historical drivers in the Amazon basin. Amazon basin contains the highest freshwater biodiversity on Earth, as so investigate phylogenetic dimension of diversity is extremally relevant from the perspective of understanding the information on the evolutionary processes that had shaped Amazonian contemporary fish assemblages.

## INTRODUCTION

Biodiversity is heterogeneously distributed among regional and local species assemblages, driven by evolutionary and ecological processes that affect rates and conditions of new species originations (e.g. speciation and dispersal), removal (e.g. extirpation and extinction), modification (e.g. adaptation) and coexistence (i.e. facilitation)(Brown, 2014; Hovikoski et al., 2007; Mittelbach et al., 2007; Ricklefs, 2007). Historical drivers often involve past geological and/or climatic events such as the formation of biogeographical barriers to species colonization (e.g., rivers and mountains formation; Rangel et al., 2018) or historical climatic variability (e.g.: Dobrovolski et al., 2012; Mascarenhas et al., 2019; Svenning et al., 2015). Evaluating such historical effects often requires the use of phylogenetic data that can provide information on the diversification history and past dispersal events that may have shaped contemporary species assemblages (Faith, 1992; Lomolino et al., 2009; K. G. Miller et al., 2005; Pigot & Etienne, 2015). For example, assemblages will be formed by closely related groups and small branch length among species if high in situ diversification is responsible for high diversity (i.e.: clustered assemblage) (Tucker et al., 2017; Webb, 2000). On the other hand, the intense dispersal of species will produce a lack of phylogenetic community structure as species would come from different branches and distinct evolutionary history (i.e., overdispersed assemblage) (Dexter et al., 2017; Tucker et al., 2017; Webb, 2000). Understanding these processes can help shed light on evolutionary community assembly and the effects of historical drivers on current distribution of organisms (Crouch et al., 2019; Dexter et al., 2017; Graham, 2003; Leprieur et al., 2016; Pyron & Burbrink, 2014; Qian et al., 2020; Brody Sandel et al., 2020).

The Amazon basin is a major biodiversity hotspot (Antonelli et al., 2018; Malhi et al., 2008), which holds the highest freshwater biodiversity on earth (Tisseuil et al., 2013). The Amazon dwelling freshwater fishes represent ~15% (>2,400 validated species) of all freshwater fish species current described worldwide (Jézéquel et al., 2020a; Tedesco et al., 2017). Despite recent advances in describing fish diversity patterns in the Amazon basin (e.g. Albert et al., 2020; Dagosta et al., 2020; Dagosta & Pinna, 2019), only one study so far has attempted to quantitatively analyze its basin-wide drivers (Oberdorff et al., 2019). Using 97 sub-drainage basins covering the entire Amazon system, this study revealed prominent influences of current climatic conditions and habitat size on sub-drainages species richness, whereas habitat size, current and past climatic stability, and isolation by waterfall better explained their endemic richness. All these drivers are already well known to promote or slow down extinction, speciation or immigration processes shaping in fine riverine fishes’ assemblage structure and diversity (Albert et al., 2020; Hugueny, B., Oberdorff, T., & Tedesco, P. A, 2010; Oberdorff et al., 2019; Tedesco et al., 2017).

More surprisingly, Oberdorff et al.(2019) also highlighted at the Amazon basin a negative upriver-downriver (West-East) gradient in species richness. This pattern is contrary to the expectation of increasing diversity at more downriver locations along fluvial systems. This reversed gradient in species richness was associated to the peculiar history of the Amazon drainage network, which, after having been isolated as western and eastern basins since the Paleogene (from ~65 Ma) (Hoorn et al., 2010), only began flowing eastward most probably during mid to late Miocene (from ~9 to 5 Ma) (Hoorn et al., 2017; Latrubesse et al., 2010). During the early Miocene (from ~23 Ma), western Amazonian was occupied by a megawetland periodically connected to the Caribbean Sea (Bicudo et al., 2019; Jaramillo et al., 2017) and subjected to multiple marine incursions (C. Hoorn et al., 2010; McDermott, 2021), known as the Pebas system (Wesselingh, 2006). This wetland system was separated from the fluvial eastern Amazonia possibly by the Purus Arch (Figueiredo et al., 2009). Following this scheme, the unexpected reverse gradient in species richness found by Oberdorff et al. (2019) suggests that the main historical center of fish diversity was located westward with a potential second center of origin located eastward, but much smaller in size and diversity and that current fish dispersal and adaptation processes from the westward center are currently progressing eastward, but not yet achieved (Oberdorff et al., 2019). However, considering the phylogenetic dimension may provide further information on the evolutionary assembly processes that may have shaped Amazonian contemporary fish species assemblages. A pattern of phylogenetic diversity congruent with the noticed reverse pattern of species richness will strengthen the hypothesis recently proposed by Fontenelle et al. (2021) that western Amazon may act as a species pump (sensu Haffer, 1969) for eastern fish assemblages, i.e.: diversity rich sub-drainages in western Amazonia due to higher speciation and persistence of older lineages, that gradually contribute lineage diversity to more species-poor sub-drainages in the East.

Further, if western Amazonia acts as a species pump for the whole Amazonian basin with currently incomplete dispersal, this would predict a pattern of geographically structured phylogenies with higher phylogenetic diversity sensu stricto for assemblages inhabiting western sub-drainages and a progressive decrease along the Amazon basin west/east gradient and with closely related species preferentially found in western Amazonia (i.e. phylogenetic clustering) due to extensive local in situ speciation in this region. Alternatively, if fish lineages have experienced frequent long-distance dispersal throughout their history, we expect random patterns of sub-drainage assemblage phylodiversity, with respect to relatedness and geography.

Here, relying on a large comprehensive database of freshwater fish species distribution in the Amazon basin (Jézéquel et al., 2020a) and taking advantage of a recently published, well-sampled molecular phylogeny of actinopterygian fishes (Rabosky, 2020; Rabosky et al., 2018), we analyzed phylogenetic diversity patterns of fish assemblages in our 97 sub-drainages. Examining the same diversity drivers as previously defined in Oberdorff et al., (2019), we determine which of these drivers were most closely associated with sub-drainage patterns of phylogenetic diversity using the phylogenetic diversity sensu stricto (PD, Faith, 1992), the mean pairwise phylogenetic distance between species in terms of branch length (MPD, Webb et al., 2002), and the mean nearest taxon distance (MNTD, Webb et al., 2002).

We first expect our three phylodiversity metrics to be, together or separately, significantly related to the same drivers already found to act significantly on sub-drainages richness and endemism patterns (Oberdorff et al., 2019). More specifically, we expect assemblages in sub-drainages that have been climatically stable overtime or located in high temperature/productivity areas to present higher phylogenetic diversity per se than expected due to persistence of phylogenetic lineages (lower extinction) and in situ diversification (i.e. positive PD and MPD, and negative MNTD values); assemblages in sub-drainages located in highly fragmented areas (i.e., with large number of waterfalls) to present lower phylogenetic diversity per se than expected due to fewer immigration events and higher phylogenetic clustering due to high in situ diversification of few lineages (i.e. negative PD, MPD, and MNTD values); assemblages in sub-drainages located in areas having suffered from past marine incursions to present lower phylogenetic diversity per se than expected due to increased lineages extinction and eventually higher phylogenetic clustering due to in situ diversification of few remaining lineages (i.e. negative PD and possibly negative MNTD values); assemblages in sub-drainages currently located in high elevation, steep gradient areas to present lower phylogenetic diversity per se than expected due to fewer immigration events and higher extinction risks in such harsh habitat conditions (i.e. negative PD and MPD values).

## MATERIAL AND METHODS

### Biological Data and Phylogeny

Occurrence records of fish have been compiled and constantly updated under the AmazonFish project (www.amazon-fish.com) by mobilizing and integrating all information available in published articles, books, gray literature, online databases, worldwide museums and Universities, and expeditions conducted during the project. The database currently contains 232,936 georeferenced records for 2,406 valid native freshwater fish species from 514 genera and 56 families (Jézéquel et al., 2020). As we were interested in riverine organisms, we excluded all species from the genus *Orestias* (i.e. 15 species) because they are mostly restricted to lakes in the Andes highlands (Scott et al., 2020). As far as we know this database contains the most complete and up-to-date information currently available on freshwater fish species distribution for the entire Amazon drainage basin.

Our defined fish assemblages consist of species presence/absence in each of the 97 sub-drainages covering the entire Amazon basin. The full methodology for sub-basins delineation is detailed in Oberdorff et al.(2019) and (Jézéquel et al., 2020). Briefly, we classified our sub-drainage basins based on the HydroBASIN framework (Lehner & Grill, 2013) and combined different HydroBASIN levels to retain only sub-basins > 20,000 km^2^ to optimize sampling effort (Oberdorff et al., 2019) (**see Appendix Fig. S1.1 in Supporting Information**).

We obtained phylogenetic information on Amazonian fishes from the supertree established by Rabosky et al. (2018). The backbone of this ultrametric supertree consists of 11,638 species for which genetic data have been available, a calibration with 130 fossil points (hereafter, genetic tree) and the insertion of 19,888 species based on a stochastic polytomy resolution algorithm (Rabosky, 2020). In the Amazon basin, we found 635 species for which genetic data were available and 1451 for which inclusion was based on the polytomy algorithm, resulting in 2086 fish species in the final pruned tree (87% of the entire fish fauna) (hereafter, genetic-polytomy tree). This final global fish tree consists of one hundred tree samples from the posterior distribution and has been consistently used to estimate fish diversification rates at the global scale (Rabosky, 2020).

### Phylogenetic fish assemblage metrics

We calculated Phylogenetic Diversity (PD), Mean Phylogenetic Diversity (MPD), and the Mean Nearest Taxon Distance (MNTD) metrics for each sub-drainage based on occurrence fish record and the phylogenetic tree data to evaluate the phylogenetic fish patterns in the Amazon basin. The first consist of the total branch length of all species occurring in a given sub-drainage (Faith, 1992), whereas the other two represent a mean relatedness of fish species (i.e., phylogenetic dispersion of clades) composing each of sub-drainage basin assemblages (Cadotte et al., 2010; Swenson, 2009; Webb, 2000). MPD describes a single assemblage by calculating mean pairwise branch distances of all species within a given assemblage, whereas MNTD calculates the mean values of only the shortest distances between taxa (Tucker et al., 2017). Although similar in computation, MPD gives information on deep evolutionary history of all taxa and broad divergent events occurring between taxa in all phylogenetic tree while MNTD focuses on divergent events occurring between taxa at the tips of the phylogenetic tree, and hence at a more recent evolutionary time scale (Webb, 2000). The use of distinct phylogenetic metrics and assemblage relatedness is justified as the historical events linked to the Amazon basin evolution occurred at different time scales (Li et al., 2019).

A potential bias related to these three phylogenetic metrics (PD, MPD and MNTD) is the fact they are all sensible to the total species richness present in each assemblage (Cadotte et al., 2010; Brody Sandel, 2018; Tucker & Cadotte, 2013). To avoid this bias, we performed a null model approach to remove the richness effects from all these metrics (hereafter named ses.PD, ses.MPD and ses.MNTD) (Kembel et al., 2010; E. T. Miller et al., 2017; Webb et al., 2002). The null model consisted in resampling with equal probability (Pigot & Etienne, 2015) 499 times the tips (i.e., species) of phylogenies and calculating all these metrics in each time step (Miller et al., 2017). We used the null distributions and the observed values to calculate the standardized effect size (ses) as (Observed – Null Simulated Mean)/Null Simulated Standard Deviation for the three metrics. For genetic+polytomy phylogeny comprising 100 phylogenetic trees sampled from the posterior distribution, we used the same null model approach for each posterior tree and calculated the mean over all 100 ses values. High positive ses values indicate high phylogenetic diversity (ses.PD) and high phylogenetic overdispersion (both with ses.MPD and ses.MNTD) in assemblages compared to the expected values of assemblages containing the same number of species. Conversely, low ses values indicate low PD and phylogenetic clustering (both with ses.MPD and ses.MNTD) compared to the null expectations based on assemblages containing the same number of species. This approach ensures our phylogenetic metrics are not affected by distinct species richness in sub-drainage basins.

Pearson correlation among metrics from only genetic and genetic+poly trees were high and statistically significant (ses.PD = 0.69, p < 0.001; ses.MPD= 0.83, p<0.001; ses.MNTD = 0.45, p<0.001). This demonstrates consistent patterns of metrics in Amazon Basin for both phylogenetic trees, so we focused here on the models using only the genetic+polytomy tree. All these metrics, null models and ses calculations have been performed under R environment (*R Core team,* 2020) using ‘pd.query’, ‘mpd.query’ and ‘mntd.query’ from *PhyloMeasures* package (Tsirogiannis & Sandel, 2016).

### Historical and contemporary drivers

Large scale biodiversity patterns can be explained by a range of ecological and historical drivers (Brown, 2014; Ricklefs, 2004), and most of these drivers also apply for freshwater fishes at large spatial scales (Hugueny, B., Oberdorff, T., & Tedesco, P. A, 2010). These drivers can be summarized under climate/productivity, area/environmental heterogeneity, historical/evolutionary, and spatial hypotheses. Data sources and definitions of the drivers used in this study are presented in detail in Oberdorff et al. (2019), and we only provide here a brief overview of each of them. All predictors described below have been extracted for each of the 97 sub-drainage basins, providing a mean single value for each of them.

We included variables related to the Amazon basin geological history from distinct time periods. We identified the sub-basins potentially belonging (1) or not (0) to the Pebas system at ~23 Mya (sensu Hoorn et al., 2010), the surface area of each sub-basin under sea water considering a sea-level rise of 25 m (<1 Mya) and of 100 m (~ 5 Mya) during recent Pleistocene marine incursions (Miller et al., 2005), and the Quaternary climate stability within the sub-basin (from ~ 21 kya to present). We used Quaternary climate reconstructions of mean, max and min annual temperatures and precipitations at the Last Glacial Maximum (LGM; 21 kyr) from three GCM models (CCSM, MIROC, and MPI) and calculated the difference between current and LGM mean values (from the three models) of the same variables to describe climate stability (Diff_CurrentLGM). The difference between the LGM climate and the present climate is one of the strongest climatic shifts in all of the Quaternary (Oberdorff et al., 2019; Sandel et al., 2011). We then performed a Principal Component Analysis (PCA) based on correlation on the difference between current-past climate and retained the first three axes explaining 88% of total variation. We considered correlation coefficients higher than 0.25 (negative or positive) as the variables better explaining each PCA axis. PC1_ Diff_CurrentLGM is positively associated to maximum precipitation (0.27), mean (0.52) and maximum (0.57) temperature; and negatively associated to minimum (−0.45) and annual precipitation (−0.32). PC2_ Diff_CurrentLGM is positively related to minimum temperature (0.29) and negatively associated with annual (0.63) and maximum (0.66) precipitation. The PC3_ Diff_CurrentLGM is positively associated to minimum precipitation (0.39) and minimum (0.87) temperature.

We further estimated the fragmentation of sub-drainage basins, a key driver of freshwater fish diversity at large scale (Dias et al., 2013), by using the number of waterfalls within each sub-basin using data available from http://wp.geog.mcgill.ca/hydrolab/hydrofalls/. We used the distance of each sub-drainages to the river mouth (km) to represent the longitudinal gradient of each of the sub-drainages within the Amazon River network (see Oberdorff et al., 2019 for a detailed explanation).

To estimate the effect of current climate and productivity, we used the annual mean and seasonality (CV of intra-year monthly values) of temperature (Temp), precipitation (Prec), actual evapotranspiration (AET), potential evapotranspiration (PET), net primary productivity (NPP), solar radiation (SolRad), run-off (RO), and the lowest (or highest) value of minimum (or maximum) temperature of the coldest (or warmest) month from WorldClim (version 1). These variables measure the mean current climatic condition, the seasonal climatic variability and the energy availability within each sub-drainage basin. We also included elevation (mean, minimum, maximum, range; in m) as climate and elevation are usually linked. Since these predictors are not the main goal here, we conducted a Principal Components Analyses (PCA) based on correlations on these variables to reduce multicollinearity. We used the first four PCA axes, which explained 85% of total variability, as synthetic predictors describing current climate and elevation gradient **(Table S1.1 in Appendix 1**). For results interpretation, we considered correlation coefficients higher than 0.25 (negative or positive) as the variables better explaining each PCA axis. PC1_GlobEnv is positively associated to net primary productivity seasonality (0.25) a negatively associated to minimum temperature (−0.26). PC2 axis is positively associated to precipitation seasonality (0.30), seasonal actual evapotranspiration (0.31), minimum (0.25) and maximum (0.37) potential net primary productivity and maximum temperature (0.30), and negatively associated to minimum precipitation (−0.29) and mean net primary productivity (−0.25). PC3_GlobEnv is positively associated to minimum solar radiation (0.29); and negatively correlated to annual precipitation (−0.27), maximum (−0.32) and annual (−0.28) actual evapotranspiration, maximum (−0.28) and seasonal (−0.30) potential evapotranspiration, and temperature seasonality (−0.34). Finally, PC4_GlobEnv is positively related to elevation: mean (0.28), maximum (0.25), variation (0.26) and range (0.37); minimum potential evapotranspiration (0.37) and mean net primary productivity (0.32); and negatively related to potential evapotranspiration seasonality (−0.29) and temperature seasonality (−0.27) **(see Table S1.1).**

Habitat size and habitat diversity were estimated using the surface area of the sub-drainage basin (km2; Area), the network density (i.e., length of the riverine network divided by the surface area of the sub-basin, a measure of habitat availability for fishes; NetwD), the land cover heterogeneity (i.e., the Shannon diversity index based on the proportion of native land cover classes within each sub-drainage basin; CoverDiv), and the soil heterogeneity (i.a., the Shannon diversity index based on the proportion of each soil type within each sub-drainage basin; SoilDiv).

Amazonian waters are divided into three distinct biogeochemical water types or “colors”, differentiated by sediment composition, geochemistry and optical characteristics: *whitewaters* have an Andean origin (e.g. the Madeira River and the Amazon mainstem), characterized by their low transparency due to large amounts of sediment particles and a neutral pH (pH ~7); nutrient poor *blackwaters* mostly draining the Precambrian Guiana shield (e.g. the Negro River) characterized by their low-transparency and acidic waters (pH < 5); and nutrient-poor, low-sediment, high-transparency, and slightly acidic *clearwaters* (pH ~ 6) mostly draining the Brazilian and Guianas shields (e.g. the Tapajós and Xingu Rivers) (Sioli, 1984). Fish assemblages structure differs between water types (e.g.: Bogotá-Gregory et al., 2020) so we classified sub-basins according to their main water type using (Venticinque et al., 2016) (**see also Fig. S2 from Oberdorff et al., 2019**). The three water types were coded as categorical variables.

Finally, the number of sampling sites divided by the surface area of each sub-basin was also included in our models (SamplingEffort). This last predictor is important to control for a potential sampling effort effect in our models as previous noticed by (Oberdorff et al., 2019).

### Statistical Analyses

Prior the analyses, we log-transformed (log[x+1]) some predictors (i.e., surface area, number of waterfalls, sampling effort, elevation mean and elevation range) to reduce the effects of extreme values. As sea level predictors are proportion values bounded between 0 and 1, we applied an arcsin square root transformation. Finally, we standardized predictors by subtracting the mean and dividing by two times the standard deviation in order to get comparable coefficients for our models (Gelman, 2008).

We fitted multiple linear regression models to determine the drivers of our three phylodiversity metrics (i.e. ses.PD, ses.MNTD and ses.MPD). We calculated the Variance Inflation Factor (VIF) for each driver after model fitting, they were all below 9, suggesting that multicollinearity was not an issue in our models (Dormann et al., 2008). We checked normality of residuals and model assumptions by drawing histograms of models’ residuals and plotting model residuals against each predictor. Using Cook’s distance, we also checked for eventual influential observations in our models.

We tested for spatial autocorrelation in model residuals by calculating the Moran’s I statistic and its p-values, using the inverse of the watercourse distance among pairs of sub-drainages as weights. When spatial autocorrelation was detected, we constructed Moran’s Eigenvector Maps (MEM, Dray et al., 2006) computed with watercourse distance. The spatial vectors related to the spatial structure for each predictor were obtained from a forward selection algorithm that avoid type I error (Blanchet et al., 2008), and we included the selected MEMs as predictors in multiple regression models to control for spatial autocorrelation. After including MEMs in models, we recalculated Variance Inflation Factor (VIF), and found high VIF values for ses.PD models for some explanatory variables (i.e.: distance, PC1_GlobEnv and MEM1, MEM3, VIF>10). However, we maintained all variables in our models to remove spatial autocorrelation.

All analyses have been conducted in R environment (*R Core Team*, 2020) using GISTools ((Chen, 2014), ggeffects (Lüdecke et al., 2021), ggplot2 (Wickham et al., 2020), vegan (Oksanen et al., 2020), rgeos (Bivand et al., 2020), effects (Fox, Weisberg, Price, Friendly, et al., 2020), adespatial (Dray et al., 2021) and car (Fox, Weisberg, Price, Adler, et al., 2020) packages.

## RESULTS

### Spatial distribution of phylogenetic diversity metrics

The ses.PD values varied between 1.88 and −4.32 (mean= −0.75, sd=1.17; **Fig. 1a**), Coari (1.88) and Blanco Baures (1.87) being the sub-drainages with highest phylogenetic diversity; Ucayali2 (−4.33) and Urubamba (−4.32) are those with the lowest values. Globally, high ses.PD values are found in sub-drainages located near the Amazon main course, and low ses.PD values in peripheral sub-drainages. Values of ses.MPD vary between 2.43 and −6.25, with mean value of −1.90 (sd= 2.15; **Fig. 1b**), suggesting that phylogenetic clustering is more frequent than overdispersion in the sub-drainages analyzed. The northeast sub-drainages Trombetas1 (2.43) and Amazon9 (1.91) have the highest positive, and the northwest sub-drainages Curaray (−6.25) and Napo2 (−5.77) the lowest ses.MPD values. As for ses.PD, we observed that ses.MPD displays the highest values along the Amazon main course and the lowest values at the periphery of the Amazon basin. For ses.MNTD, more negative than positive values are also found (mean= −1.03, sd= 0.96; range= −3.98−0.58; **Fig. 1c**), suggesting that most sub-drainages present a clustered phylogenetic pattern. The sub-drainages Maues (0.58) and Jamanxim (0.58) have the largest positive ses.MNTD values, and Urubamba (−3.98) and Mantaro (−3.20) the highest negative values.

**Fig. 1.**
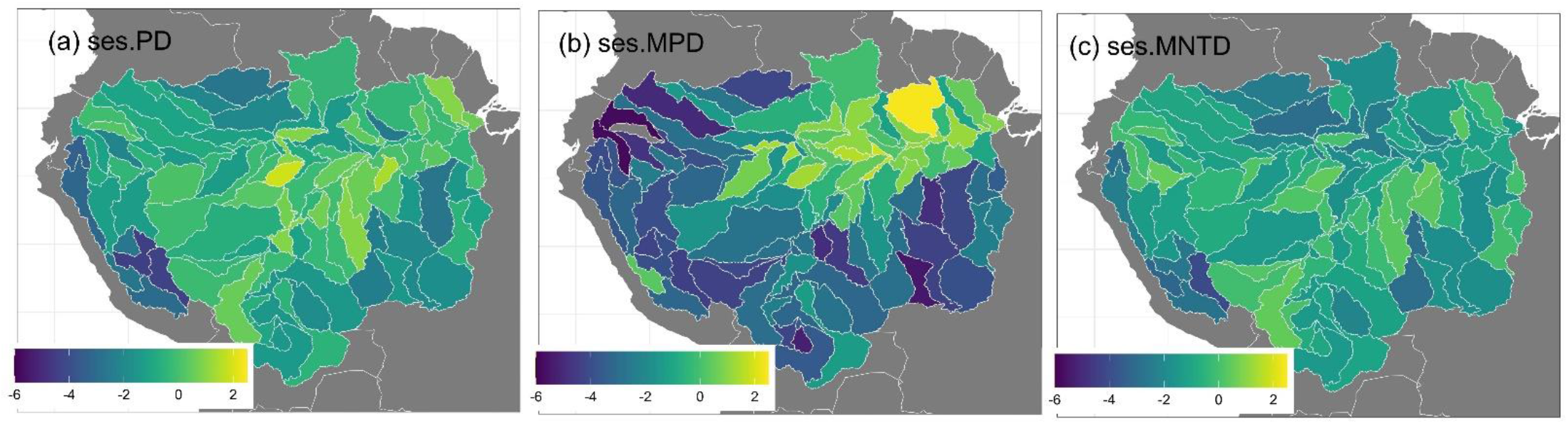
Standardized effect size of phylogenetic diversity calculated using 2086 native Amazonian freshwater fish species (genetic+polytomy tree). (a) sensu stricto (ses.PD), (b) mean phylogenetic pairwise distance (ses.MPD), and (c) mean nearest taxon distance (ses.MNTD). Negative values of ses.MNTD and ses.MPD indicate phylogenetically clustered assemblages, while positive ones phylogenetically overdispersed assemblages.

Phylogenetic metrics calculated from genetic tree (635 fish species) presented a similar distribution to the three phylogenetic metrics calculated from genetic+polytomy tree (**see Fig. S2.6 in Supporting Information**). The ses.PD metric varies between 3.52 and −2.28 (0.51, sd=1.08; **Figure S2.6a**). We found ses.MPD more negative values varying between 1.39 and −5.28 (mean=−1.58, sd= 1.88; **Figure S2.6b**) and ses.MNTD varying between 3.07 and −2.35 (−0.27, sd=1.09; **Figure S2.6c**).

Pearson correlations show that ses.PD, ses.MPD and ses.MNTD are unrelated to taxonomic metrics (richness and endemism), whereas ses.PD is correlated to ses.MPD and ses.MNTD (lower triangle, **Table 1;** see **Table S2.2 in Supporting Information** for genetic tree results). The raw values of all phylogenetic metrics showed much high correlation values among each other and the other taxonomic metrics (upper triangle, **Table 1;** see **Table S2.2** for genetic tree results).

**Table 1.**
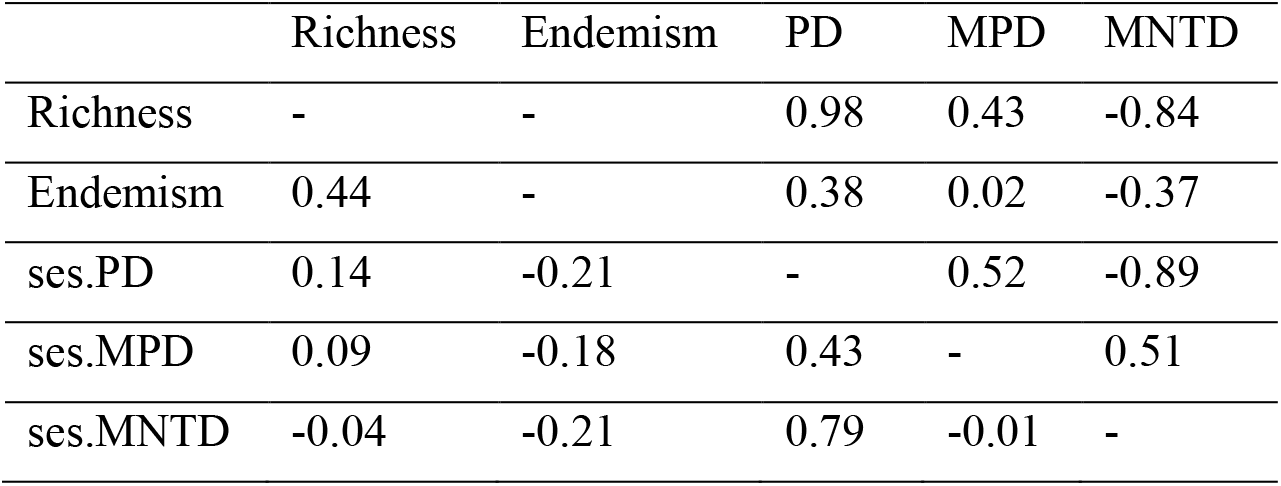
Pearson Correlation coefficients between species richness, endemism, and each of the three phylogenetic metrics calculated from the genetic+polytomy tree for 2086 freshwater fish species (phylogenetic diversity, PD; mean pairwise distance, MPD; and mean nearest taxa distance, MNTD). Values above the diagonal (upper triangle) refer to correlations between sub-drainages total species richness, endemic richness and raw phylogenetic metric values (i.e. without controlling for richness). Values below (lower triangle) refer to correlations between sub-drainages total species richness, endemic richness and standardized effect size (ses.) of the three phylogenetic metrics.

|  | Richness | Endemism | PD | MPD | MNTD |
| --- | --- | --- | --- | --- | --- |
| Richness | - | - | 0.98 | 0.43 | -0.84 |
| Endemism | 0.44 | - | 0.38 | 0.02 | -0.37 |
| ses.PD | 0.14 | -0.21 | - | 0.52 | -0.89 |
| ses.MPD | 0.09 | -0.18 | 0.43 | - | 0.51 |
| ses.MNTD | -0.04 | -0.21 | 0.79 | -0.01 | - |

### Determinants of the three phylogenetic diversity metrics

None of our selected historical predictors have effect on ses.PD, this last metric being only significantly negatively related to sub-drainage surface area and distance of the sub-drainage from the Amazon River mouth (**Fig. 2a**; **Table 2**). Note that there is also a marginally significant positive effect of sub-drainage network density (i.e. the length of the riverine network divided by the surface area of the sub-drainage) on ses.PD. There is a significant spatial structure in model residuals, which has been controlled for by including the three selected MEMs (**Fig. S1.2**).

**Table 2.** Estimates, 95% confidence interval and p-values from the multiple regression models for phylogenetic diversity estimated using ses.PD, ses.MPD and ses.MNTD. Significant relationships (p<0.05) are in bold.

|  | ses.PD |  | ses.MPD |  | ses.MNTD |  |
| --- | --- | --- | --- | --- | --- | --- |
|  | *Estimates (CI)* | *p* | *Estimates (CI)* | *p* | *Estimates (CI)* | *p* |
| (Intercept) | **-1.05 (-1.95 – -0.15)** | **0.023** | -1.00 (-2.22 – 0.22) | 0.108 | **-1.27 (-2.09 – -0.45)** | **0.003** |
| WaterColor [Clear] | -0.20 (-1.14 – 0.75) | 0.683 | -1.10 (-2.35 – 0.16) | 0.085 | -0.24 (-1.07 – 0.60) | 0.579 |
| WaterColor [White] | 0.48 (-0.31 – 1.27) | 0.229 | **-1.18 (-2.28 – -0.08)** | **0.036** | 0.28 (-0.45 – 1.02) | 0.449 |
| NetwD | 0.19 (-0.02 – 0.40) | 0.075 | -0.12 (-0.42 – 0.18) | 0.432 | 0.12 (-0.08 – 0.32) | 0.230 |
| Area | **-0.33 (-0.63 – -0.03)** | **0.030** | 0.02 (-0.35 – 0.39) | 0.909 | **-0.27 (-0.51 – -0.02)** | **0.033** |
| SoilDiv | 0.08 (-0.17 – 0.33) | 0.506 | 0.24 (-0.10 – 0.58) | 0.160 | 0.05 (-0.18 – 0.27) | 0.686 |
| DistMouth | **-1.29 (-2.12 – -0.46)** | **0.003** | **-0.99 (-1.73 – -0.25)** | **0.009** | **-0.61 (-1.11 – -0.12)** | **0.016** |
| CoverDiv | -0.12 (-0.47 – 0.23) | 0.511 | -0.01 (-0.50 – 0.48) | 0.959 | -0.01 (-0.34 – 0.32) | 0.945 |
| PC1_Diff_CurrentLGM | -0.08 (-0.45 – 0.29) | 0.654 | 0.37 (-0.14 – 0.87) | 0.151 | 0.14 (-0.20 – 0.48) | 0.413 |
| PC2_Diff_CurrentLGM | -0.01 (-0.36 – 0.34) | 0.959 | -0.22 (-0.66 – 0.21) | 0.315 | 0.01 (-0.28 – 0.31) | 0.920 |
| PC3_Diff_CurrentLGM | -0.04 (-0.35 – 0.27) | 0.803 | -0.19 (-0.61 – 0.23) | 0.379 | 0.07 (-0.21 – 0.35) | 0.615 |
| PebasLake | 0.08 (-0.72 – 0.88) | 0.844 | 0.17 (-0.82 – 1.16) | 0.726 | 0.20 (-0.46 – 0.86) | 0.551 |
| Sea water at <1Mya | -0.26 (-0.63 – 0.11) | 0.165 | 0.42 (-0.10 – 0.93) | 0.111 | **-0.35 (-0.69 – 0.00)** | **0.050** |
| Sea water at ~5Mya | 0.16 (-0.19 – 0.51) | 0.358 | **0.71 (0.22 – 1.20)** | **0.005** | -0.14 (-0.47 – 0.19) | 0.397 |
| Waterfall | -0.16 (-0.42 – 0.09) | 0.201 | 0.20 (-0.14 – 0.53) | 0.240 | **-0.29 (-0.51 – -0.06)** | **0.012** |
| PC1_GlobEnv | 0.46 (-0.19 – 1.11) | 0.162 | -0.07 (-0.82 – 0.67) | 0.847 | -0.15 (-0.65 – 0.35) | 0.548 |
| PC2_GlobEnv | -0.17 (-0.61 – 0.27) | 0.449 | 0.45 (-0.08 – 0.98) | 0.093 | 0.04 (-0.32 – 0.39) | 0.827 |
| PC3_GlobEnv | -0.17 (-0.46 – 0.11) | 0.225 | **0.55 (0.19 – 0.92)** | **0.003** | -0.23 (-0.48 – 0.01) | 0.063 |
| PC4_GlobEnv | -0.14 (-0.38 – 0.11) | 0.268 | **-0.43 (-0.75 – -0.10)** | **0.011** | -0.00 (-0.22 – 0.22) | 0.989 |
| SamplingEffort | -0.22 (-0.48 – 0.04) | 0.096 | -0.22 (-0.58 – 0.15) | 0.242 | -0.22 (-0.46 – 0.03) | 0.084 |
| MEM3 | **-0.47 (-0.84 – -0.11)** | **0.012** | - | - | - | - |
| MEM1 | 0.18 (-0.40 – 0.77) | 0.537 | - | - | - | - |
| MEM6 | -0.16 (-0.36 – 0.05) | 0.127 | - | - | - | - |
| R<sup>2</sup> / R<sup>2</sup> adjusted | 0.587 / 0.464 |  | 0.738 / 0.674 |  | 0.414 / 0.270 |  |
| Moran's I (*p* value) | -0.01 (*p*= 0.67) |  | -0.01 (*p*= 0.60) |  | -0.01 (*p*= 0.14) |  |

**Fig. 2.**
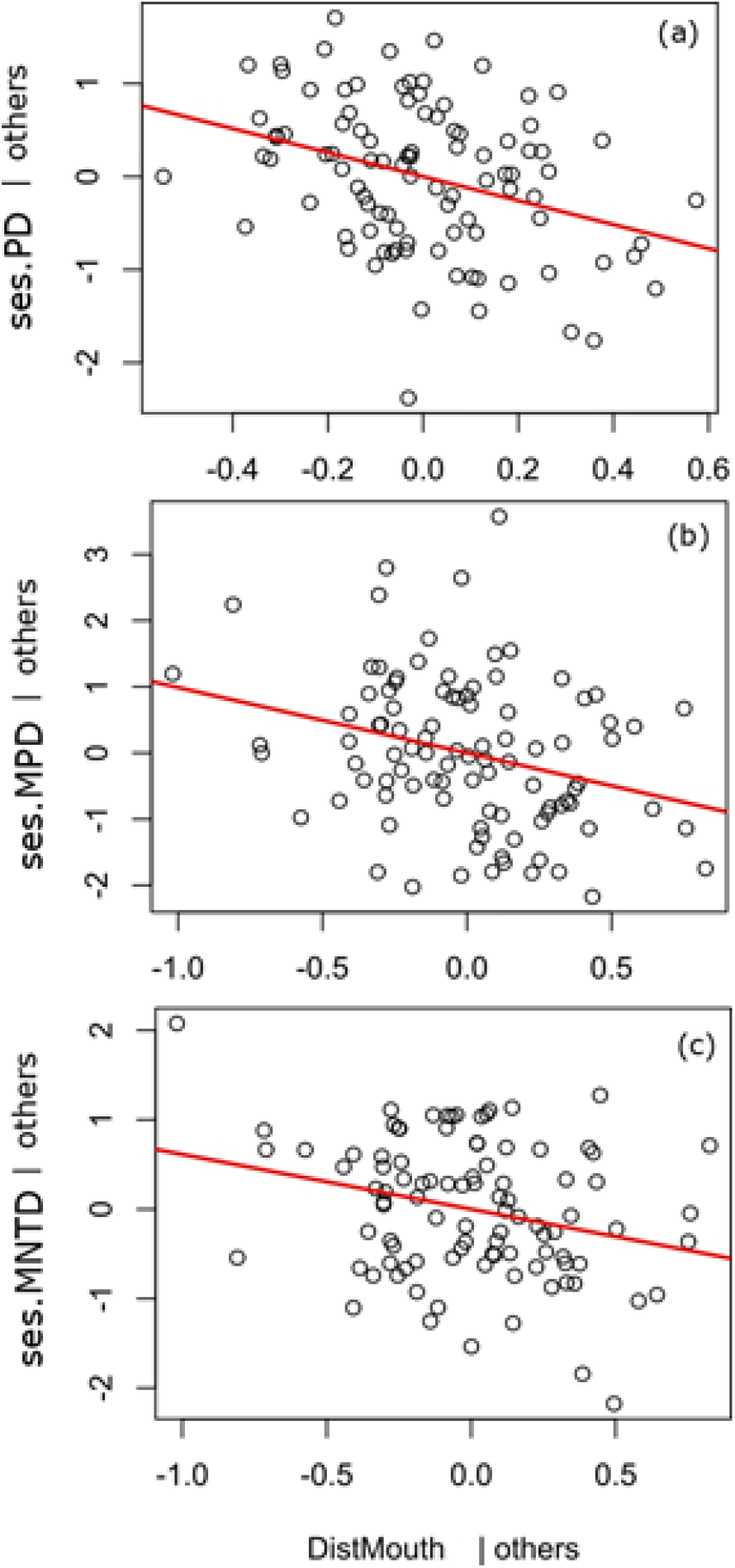
Significant relationships between phylogenetic diversity (ses.PD) and phylogenetic relatedness (ses.MNTD and ses.MPD) metrics and the distance of sub-drainages from the river mouth, after controlling for all other drivers considered in our model. Axes represent partial regression residuals controlling for all other predictors and extracted from multiple regression models; these effects are calculated considering mean values of all the other predictors as they are scaled so that their mean values equal zero.

The ses.MPD is positively influenced by the surface of sub-drainage covered by sea water at ~5Mya (**Fig. 3b**), negatively related to distance from river mouth (**Fig. 2b**) and related to current climate (positive effect of PC3_globEnv, and negatively related to PC4_GlobEnv) (**Fig. S1.3-S1.4**; **Table S1.1**). Values of ses.MPD are also significantly related to water types, white and clear waters having lower values compared to black waters (**Fig. S1.5**). Finally, significant negative effects of natural fragmentation (i.e. the total number of waterfall; **Fig. 3a**), sub-drainage surface covered by sea water at <1Mya (**Fig. 3c**), distance from river mouth (**Fig. 2c**), and surface area of sub-drainage are observed for ses.MNTD values. Similar coefficients emerged when evaluating the same models with genetic tree even if significance of predictors changed slightly (**see Table S2.3**).

**Fig. 3.**
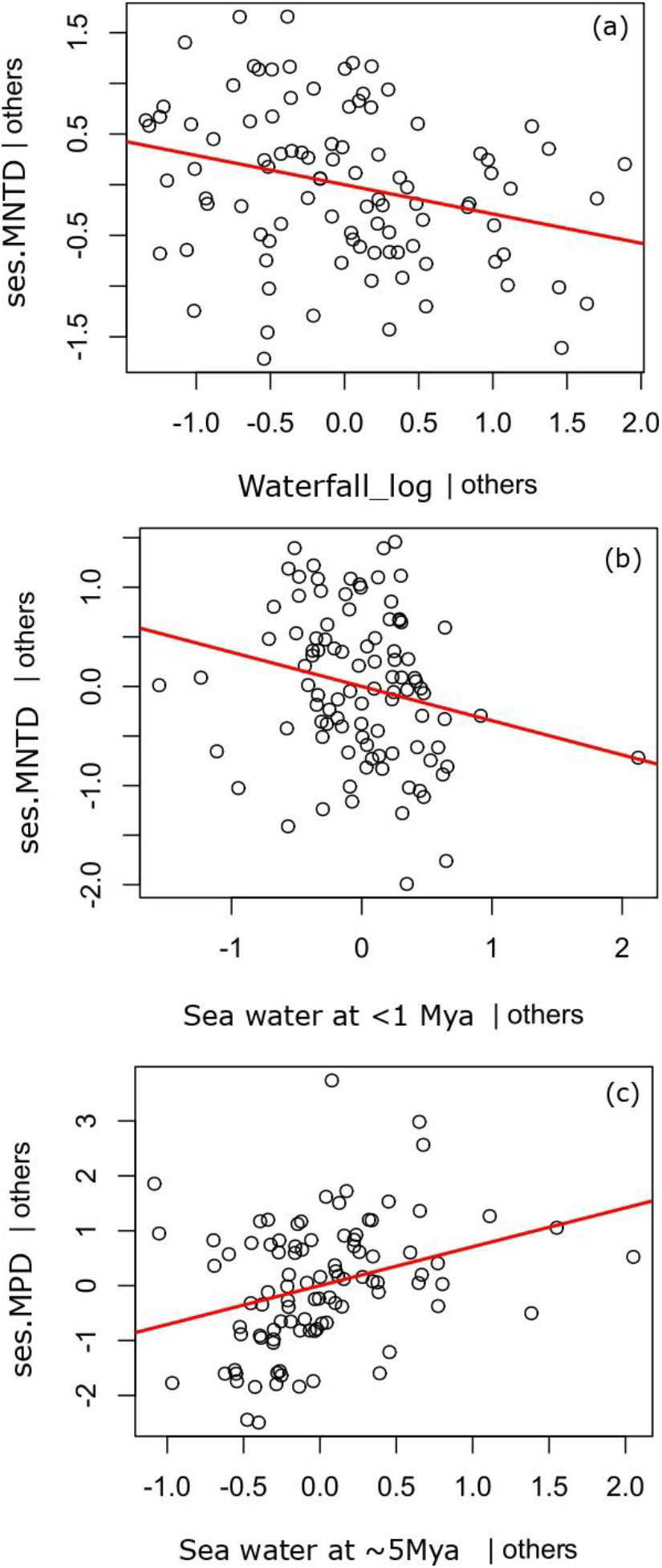
Significant relationships between phylogenetic relatedness metrics (ses.MNTD and ses.MPD) and (a) Habitat natural fragmentation (Waterfall_log), (b) Proportion of sub-drainages surface covered by seawater at <1Mya (25m) and (c) Proportion of sub-drainages surface covered by seawater at ~5Mya (100m), after controlling for all other drivers considered in our model. These effects are partial regression residuals (see Fig 2 for further details).

## DISCUSSION

Both contemporary and historical drivers play an important role in explaining current patterns of biodiversity (Coronado et al., 2015; Crouch et al., 2019; Ricklefs, 1987, 2004). Here, using a large data set on freshwater fish distribution in the Amazon basin (Jézéquel et al., 2020) and a recent phylogenetic super tree (Rabosky, 2020), we found marked and contrasting phylogenetic signatures of fish assemblages across the 97 sub-drainages, whatever the phylogenetic metric analyzed (i.e. ses.PD, ses.MPD, ses.MNTD). As far as we know, this is the first study applying this phylogenetic framework and its drivers at the scale of the whole Amazon basin (but see e.g.: Craig et al., 2020 for an application at more regional scale and Dagosta et al., 2021). These contrasting signatures were related to various historical and contemporary drivers most often similar, as expected, to the ones related to richness and endemism patterns (Oberdorff et al., 2019).

### Regional scale contemporary drivers of Amazonian phylodiversity patterns

We found effects of some current environmental drivers on assemblage phylogenetic structure in our 97 sub-drainages. The third and fourth PCA axes describing current climate were respectively positively and negatively linked to ses.MPD values that captures the variation in phylodiversity at deep time scales. The former is linked to high energy availability and stable climatic conditions whereas the latter is negatively related to high elevation and steep gradients (**see Fig. S1.3 and S1.4 in Appendix 1**). Both relationships indicate more overdispersed fish assemblages in sub-drainages located in the eastern part along the Amazon mainstem, in sub-drainages located in low elevation and in less steep gradient areas. Sub-drainages in such conditions may accumulate lineages due to lower extinction and higher colonization probabilities (Carvajal-Quintero et al., 2019; Oberdorff et al., 2019). High elevation areas, on the other hand, are more restricted to colonization by highly adapted species and subject to numerous random extinction events due to habitat harshness (Datry et al., 2016). This may result in species poor assemblages with low lineage diversification over the long term. High fish endemism levels in stable climatic sub-drainages and low total richness in sub-drainages with high elevation and steep gradients are reported in Oberdorff et al. (2019), but our results point only to a marginally significant and negative relationship between ses.MNTD and the stability PCA axis.

Our results also depict an effect of water types on ses.MPD, black waters hosting assemblages more phylogenetically overdispersed (only at high phylogenetic levels) than their clear and white waters counterparts (**Fig. S1.5**). This pattern may be due to the specific characteristics of black water stained by tannins and humic acids leached from vegetation and having low pH (pH~5 or lower) and low autochthonous productivity (Bogotá-Gregory et al., 2020). These characteristics create strong barriers to colonization for species unfitting these specific conditions that necessitate long term adaptation (Beheregaray et al., 2015; Crampton, 2019; Dagosta & Pinna, 2019; Gonzalez, Wilson & Wood, 2006; Van Nynatten et al., 2015) and may have thus promoted lineages diversification through isolation processes.

### Regional scale historical drivers of Amazonian phylodiversity patterns

Freshwater fishes are highly limited by connectivity among habitats (Carvajal-Quintero et al., 2019; Rahel, 2007; Tonkin et al., 2018). The negative relationship found between the number of waterfalls and the phylogenetic metric ses.MNTD (**Fig. 3a**), capturing the variation in phylodiversity at shallow evolutionary time, suggests that intensely fragmented sub-drainages have assemblages formed by closely related species (i.e., assemblage showing a phylogenetic clustering pattern), which can be attributed to allopatric speciation due to reduced population dispersal and consequent reduced gene flow among fragmented populations (Dias et al., 2013; Tedesco et al., 2012). Peripheral sub-drainages of the Amazon basin drain fragmented landscapes (e.g.: ancient, cratonic rivers in the Brazilian and Guiana shields and recent Andean mountains) (Bicudo et al., 2019; Hoorn et al., 2010) and contain high fish endemism levels (Oberdorff et al., 2019). Furthermore, although ses.PD and ses.MPD showed no significant link with habitat fragmentation by waterfalls, low associated values of both metrics indicate lower phylogenetic diversity sensu stricto than expected in sub-drainages with comparable species richness. Together, these findings support the idea of recent speciation events coupled with both high extinction and/or low colonization rates in fragmented sub-drainages (Albert et al., 2011).

The Amazon basin has been subject in the past to a series of marine incursions in both its western (e.g.: Bicudo et al., 2019; C. Hoorn et al., 2010) and eastern (e.g.: Christ et al., 2021) parts and on ancient (Eocene to Miocene periods in the West; Pozo and Pebas systems) and more recent (Pleistocene marine incursions in the East) time scales. These marine incursions have favored the adaptation of several marine lineages to fresh water environment (Bloom & Lovejoy, 2017; Fontenelle et al., 2021; Lovejoy et al., 2006) but also probably led to high extinctions due to the concomitant elimination of freshwater habitats in areas affected by these marine incursions (Oberdorff et al., 2019). In agreement with this last hypothesis, we found a slightly significant negative relationship between ses.MNTD and eastern sub-drainages impacted by the last see level rise during the middle Pleistocene (<1Mya, ~25m) (Christ et al., 2021), meaning that assemblages in these sub-drainages are more phylogenetically clustered than expected. This may be due to high extinction rates of lowland freshwater fish species in submerged areas that reduced their overall species richness and diversification processes in the remaining, higher elevation, isolated areas (Oberdorff et al. 2019). We also found that the phylogenetic metric ses.MPD, capturing the variation in phylodiversity at high phylogenetic levels, was positively related to eastern sub-drainages having experienced older and longer marine incursions (~5Mya, from 50 to 100 m and a duration of ~0.8 Mya) (Haq et al., 1987). Given that phylogenetic diversity sensu stricto (ses.PD) is not significantly higher than in other sub-drainages and that many marine-derived species such as anchovies, flatfishes, pufferfishes, drum, needlefishes, and stingrays are present in these sub-drainages, we suggest that the increase in ses.MPD is related to the presence of marine-derived lineages in these sub-drainage assemblages.

The presence, extension, and attributes of the Pebas system is still uncertain (Bicudo et al., 2019; Hoorn et al., 2010; McDermott, 2021), and our results show no significant effect of our categorical variable the “Pebas Lake system” (sensu Hoorn et al., 2010) on assemblage phylodiversity patterns. There was no marked indication of structured phylogenetic diversity – overdispersed or clustered fish assemblages structured phylogenies – that could have been produced by extinction, dispersal and diversification of lineages within the Pebas system. However, the rather rough variable we used here may have missed some key areas of the historical Pebas system, failing to capture any significant phylogenetic structure. This is most probable as we did find a strong phylogenetic structure along the Amazonian West/East gradient. As previously said, the geographical boundaries of the Pebas are uncertain and vary substantially according to authors (Fontenelle et al., 2021).

### Basin wide drivers of phylodiversity

We found a significant pattern of fish assemblage phylogenetic diversity along the Amazon west/east gradient. Phylogenetic diversity sensu stricto (ses.PD) and ses.MPD metric increases in sub-drainages from west to east meaning that western assemblages are less phylogenetically diversified, at least in term of lineages, than eastern ones (**Fig. 1a, b**). This result is inconsistent with the prediction of higher lineages diversity in sub-drainages of Western Amazonia and thus refutes the hypothesis that this region acts as a broad species pump for the whole basin, as recently suggested by Fontenelle et al. (2021) for marine-derived lineages only. This increase in assemblages phylogenetic diversity (sensu stricto) from west to east, already highlighted by Dagosta et al. (2020), is surprising insofar as this pattern is also opposite to the one noticed by Oberdorff et al. (2019) for species richness (i.e. sub-drainages species richness slightly but significantly decreases along a West-East gradient). However, we also found here a decreasing pattern of phylogenetic clustering from west to east (as measured by the ses.MNTD metric, **Fig. 1c**) suggesting large and recent radiations of fewer lineages in sub-drainages of the western Amazon that may have generated in fine higher overall species richness over time compared to more eastern ones. Following this pattern, western Amazonia can be seen as an evolutionary cradle of biodiversity (i.e.: location with unusually high rates of speciation) rather than a species pump region. On the other hand, the lower phylogenetic diversity in western assemblages compared to eastern ones suggests either historical unsuccessful colonization events or most probably intense lineages extinction in this region.

Indeed, the repeated transitions between Eocene and Miocene periods from fluvial-like systems to wetlands (Pozo and Pebas systems)(Antoine et al., 2016; Bicudo et al., 2019) produced strong habitat filtering for species that, together with complex salinity gradients due to periodical connections of the system to the Caribbean region, may have promoted some lineages extinction and remaining lineages diversification in the successive fluvio/lacustrine systems. The analysis of fossil records potentially available in this region (e.g.: Chabain et al., 2017) may shed further light on these possible extinction processes (e.g.: Chabain et al., 2017). In contrast, the easterly flowing proto-Amazon system (eastward of the Purus Arch) has been geologically and hydrologically more stable than western Amazonia during the past 30 millions of years (Bicudo et al., 2019; Hoorn et al., 2010; Sacek, 2014), probably causing higher phylogenetic diversity sensu stricto due to accumulation and persistence of lineages in this area over a longer period of time (Coronado et al., 2015). Furthermore, fish assemblages in this eastern region (downstream part of the Amazon current longitudinal gradient and near the historical West-East Amazon divide of Purus Arch) may benefit from lineages accumulation due to the conjunction of the three Amazon water types and to local colonization of species historically inhabiting both sides of the historical barrier (Dagosta et al., 2020).

To conclude, our study has revealed a highly non-random spatial and environmental distribution of our three phylogenetic diversity metrics (ses.PD, ses.MPD, ses.MNTD) across the 97 sub-drainages covering the Amazon basin. This suggests that diversification most often occurs within specific geographic areas (e.g.: fragmented areas, water types, western vs eastern Amazonia) and that the long-distance dispersal of species is still rare among these areas (but see Fontenelle et al. 2021 for marine-derived lineages). However, it is still relevant analyze the variation in species composition between our 97 subdrainage assemblages (i.e., taxonomic Beta diversity, sensu Whittaker, 1960), since it will bring a more precise picture of the effect of dispersal limitation in shaping current fish assemblages in the Amazon basin.

## Data availability statement

Species present in each sub-drainage (Jézéquel et al. 2020), global scale phylogeny (Rabosky et al. 2018, Rabosky 2020), and predictors used in this study (Oberdorff et al. 2019) are available in their corresponding citation. Further phylogenetic values and its standardized effect size (ses.) will be available after final acceptance and also upon request to authors.

## Acknowledgement

This paper is dedicated to J. Maldonado-Ocampo, who recently passed away during a fishing trip in the Río Vaupés (Colombia). This research benefited from support from the ERANet-LAC (www.eranet-lac.eu/) “AmazonFish” (ELAC2014/ DCC−0210) project. We thank the Field Museum of Natural History in Chicago and the Jhon D. and Catherine T. MacArthur Foundation (Grant G-1607-151047) for having mobilized data for AmazonFish in Colombia, Ecuador, and Peru. French Laboratories of Excellence “CEBA” (ANR-10-LABX-25-01) and “TULIP” (ANR-10-LABX-41 and ANR-11-IDEX-0002-02) are also acknowledged. M.S.D. thanks CNPq (150784/2015-5) and Fundação de Amparo à Pesquisa do Distrito Federal (FAPDF, #00193.00001819/2018-75, #00193.00000002/2019-61) for funding and TO, BH and PAT thanks the CEBA project “EMERGENCE”. All data were collected through the AmazonFish project (www.amazon-fish.com).

## Biosketch

Laís Salgueiro is a freshwater ecologist mostly interested in the study of biogeography and phylogenetic patterns in freshwater organisms.

## Author contributions

MSD, LS, and TO conceived the study. MSD and CJ performed GIS analyses, LS performed statistical analyses with MSD and TO advices. LS, MSD, FC and TO wrote the first draft, and all coauthors assisted in writing and revising the final version of the manuscript.

