## Supporting Information for "Drivers of phylogenetic structure in Amazonian freshwater fish assemblages"

### Appendix 1

**Table S1.1.** Loadings for each climatic (CurrClimate) and elevation (Elev) variable on the four PCA axes retained. Elev = Elevation. Temp = Temperature. Prec = Precipitation. SolRad and RO = Solar radiation and run off. AET, PET, and NPP = Actual Evapotranspiration, Potential Evapotranspiration, and Net Primary Productivity; Min, Max, Mean, Annual, CV, Range, and SD = minimum, maximum, mean, total annual, Coefficient of Variation, range, and Standard Deviation.

| | **PC1** | **PC2** | **PC3** | **PC4** |
|---|---|---|---|---|
| ElevMean_log | 0.24 | -0.13 | -0.10 | **0.28** |
| Elev_max | 0.20 | -0.23 | -0.15 | **0.25** |
| Elev_min | 0.23 | -0.08 | -0.02 | -0.07 |
| Elev_std | 0.19 | -0.22 | -0.15 | **0.26** |
| ElevRge_log | 0.20 | -0.13 | -0.20 | **0.37** |
| CurrClimate_precAnn | -0.23 | -0.12 | **-0.27** | 0.04 |
| CurrClimate_precMin | -0.15 | **-0.29** | -0.21 | -0.16 |
| CurrClimate_precCV | 0.16 | **0.30** | 0.16 | 0.15 |
| CurrClimate_precMax | -0.18 | 0.11 | -0.23 | 0.24 |
| CurrClimate_ro_mean | -0.21 | -0.06 | -0.04 | 0.08 |
| CurrClimate_aetMax | -0.15 | 0.20 | **-0.32** | 0.17 |
| CurrClimate_aetMin | -0.20 | -0.24 | -0.19 | -0.17 |
| CurrClimate_aetAnn | -0.24 | -0.08 | **-0.28** | 0.03 |
| CurrClimate_aetCV | 0.18 | **0.31** | 0.05 | 0.16 |
| CurrClimate_petMax | 0.01 | **0.37** | **-0.28** | 0.09 |
| CurrClimate_petMin | -0.17 | **0.25** | 0.10 | **0.37** |
| CurrClimate_petCV | 0.21 | 0.10 | **-0.30** | **-0.29** |
| CurrClimate_npp_mn | -0.12 | **-0.25** | -0.12 | **0.32** |
| CurrClimate_npp_cv | **0.25** | -0.08 | -0.12 | 0.01 |
| CurrClimate_tempMin | **-0.26** | 0.07 | -0.03 | -0.05 |
| CurrClimate_tempMax | -0.17 | **0.30** | -0.18 | 0.02 |
| CurrClimate_tempMean | -0.24 | 0.18 | -0.14 | -0.04 |
| CurrClimate_tempCV | 0.21 | 0.08 | **-0.34** | **-0.27** |
| CurrClimate_SolRad_mn | -0.22 | -0.14 | **0.29** | 0.18 |
| CurrClimate_SolRad_cv | 0.20 | 0.17 | -0.18 | 0.00 |
| **Proportion of variance** | 0.49 | 0.23 | 0.08 | 0.05 |
| **Cumulative proportion** | 0.49 | 0.72 | 0.80 | 0.85 |

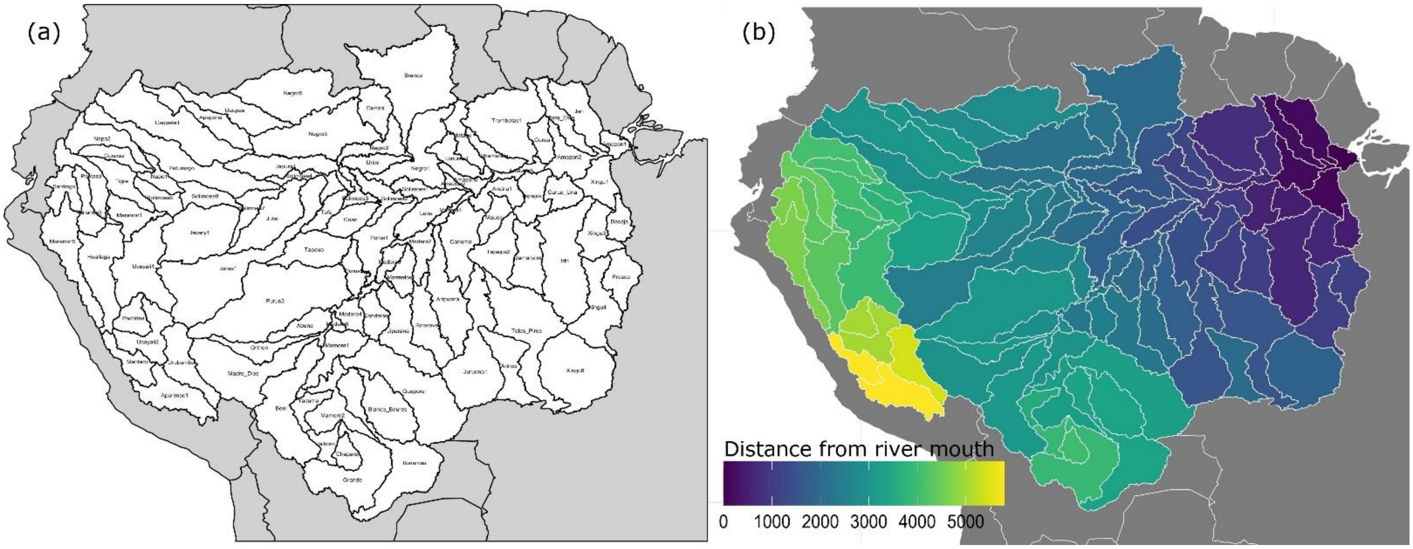


**Fig. S1.1.** (a) Names and localization of the 97 sub-drainages and (b) distance (km) of each sub-drainage from the Amazon river mouth (from dark blue near the mouth to yellow for most upstream sub-drainages).

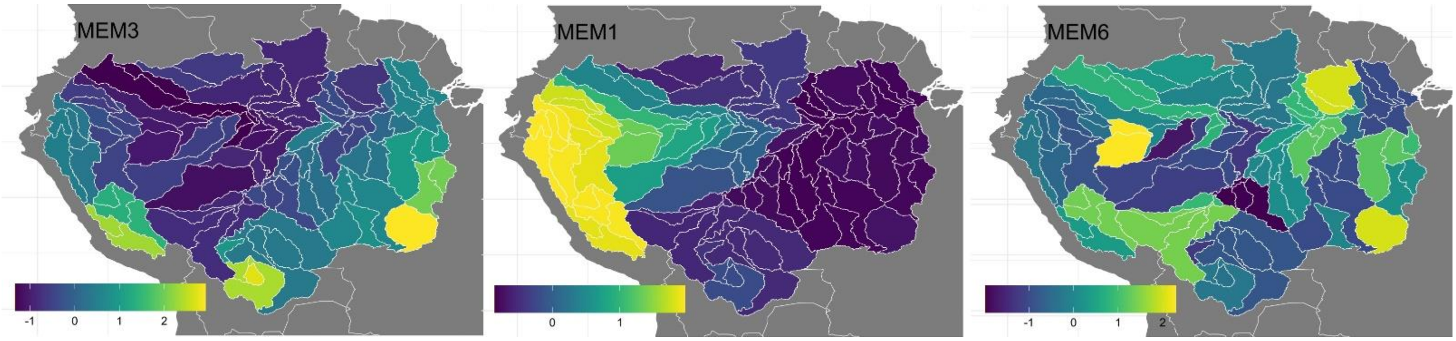


**Fig. S1.2.** Selected axis of Moran Eigenvector Maps (MEM) included in the ses.PD model. This spatial structure has been calculated based on watercourse spatial distance between all sub-drainage basins.

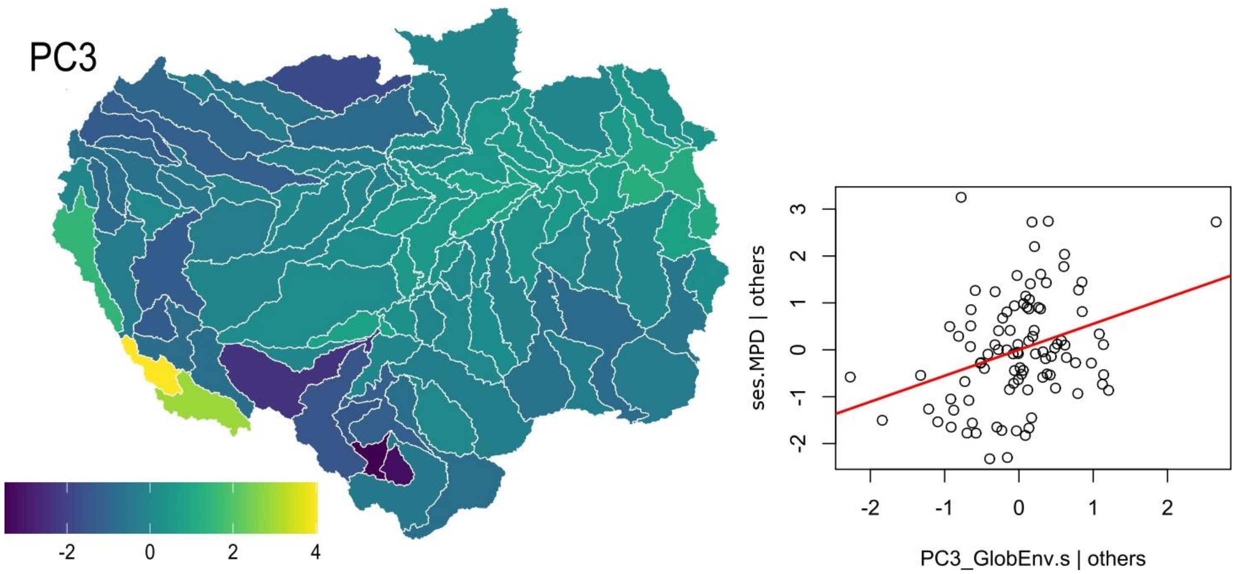


**Fig. S1.3.** Distribution values of PCA axis 3 (PC3_globEnv) for our 97 sub-drainages and its significant relationship with the phylogenetic metric ses.MPD (calculated for 2086 Amazonian freshwater fish species with genetic+polytomy tree), after controlling for all other drivers considered in our model.

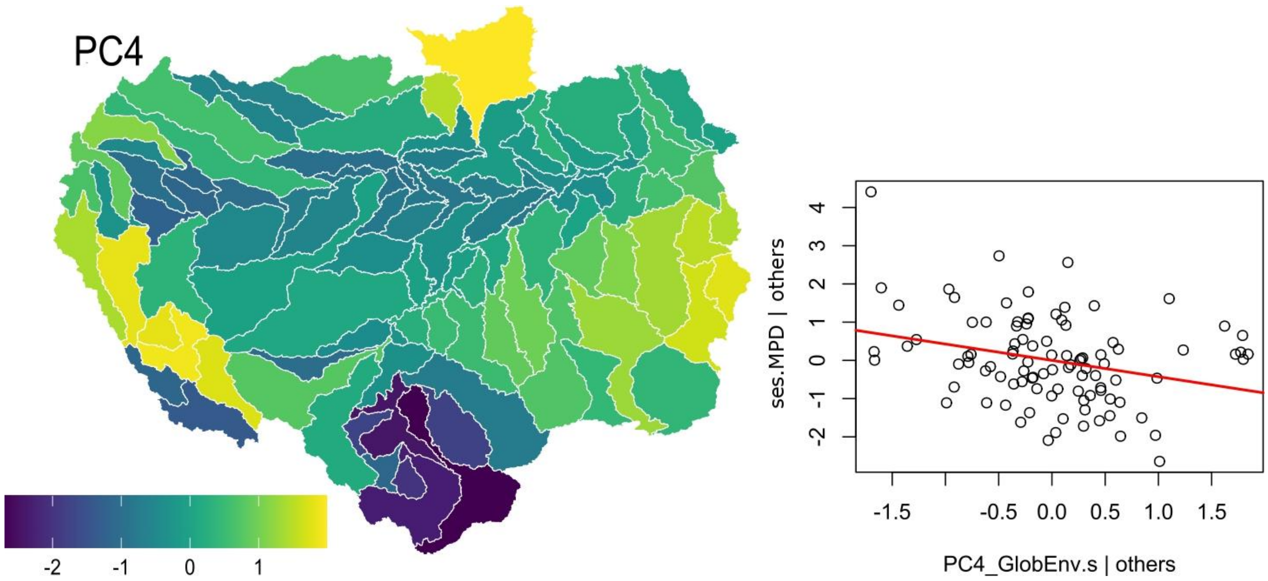


**Fig. S1.4.** Distribution values of PCA axis 4 (PC4_globEnv) for our 97 sub-drainages and its significant relationship with the phylogenetic metric ses.MPD (calculated for 2086 Amazonian freshwater fish species with genetic+polytomy tree), after controlling for all other drivers considered in our model.

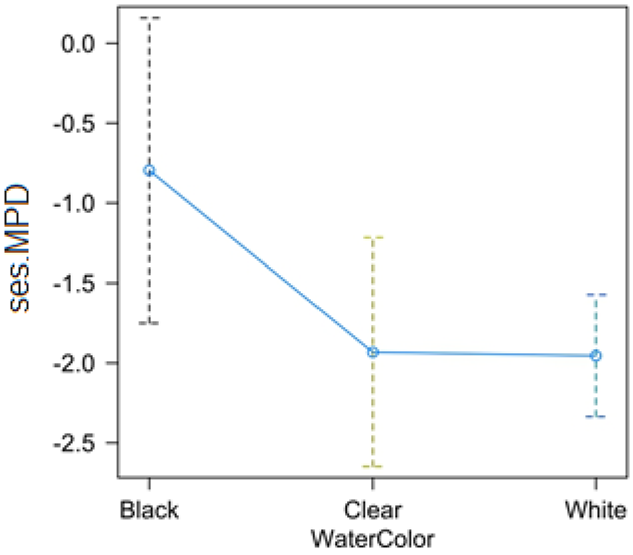


**Fig. S1.5.** Predicted mean values showing a significant effect of water type on ses.MPD values (calculated for 2086 Amazonian freshwater fish species with genetic+polytomy tree).

**Appendix 2**

**Table S2.2**. Pearson Correlation coefficients between species richness, endemism, and each of the three phylogenetic metrics calculated from the genetic tree of 635 freshwater fish species (phylogenetic diversity, PD; mean pairwise distance, MPD; and mean nearest taxa distance, MNTD). Values above the diagonal (upper triangle) refer to correlations between sub-drainages total species richness, endemic richness and raw phylogenetic metric values (i.e. without controlling for richness). Values below (lower triangle) refer to correlations between sub-drainages total species richness, endemic richness and standardized effect size (ses.) of the three phylogenetic metrics.

| | Richness | Endemism | PD | MPD | MNTD |
|---|---|---|---|---|---|
| Richness | - | 0.44 | **0.92** | 0.28 | **-0.67** |
| Endemism | **0.44** | - | 0.31 | 0.05 | -0.24 |
| ses.PD | 0.11 | -0.05 | - | 0.40 | **-0.69** |
| ses.MPD | -0.03 | -0.14 | **0.47** | - | 0.19 |
| ses.MNTD | -0.37 | -0.20 | **0.76** | 0.33 | - |

**Table S2.3**. Estimates, confidence intervals and p-values of the Regression models for the phylogenetic relatedness estimated with MNTD, MPD and PD metrics for 635 Amazon freshwater fish species in 97 sub-drainages, obtained by the genetic tree.

| | **ses.PD_gen** | | **ses.MPD_gen** | | **ses.MNTD_gen** | |
|---|---|---|---|---|---|---|
| | *Estimates (CI)* | *p* | *Estimates (CI)* | *p* | *Estimates (CI)* | *p* |
| (Intercept) | 0.31 (-0.63 – 1.24) | 0.517 | -0.94 (-1.99 – 0.12) | 0.081 | 0.08 (-0.85 – 1.00) | 0.871 |
| WaterColor [Clear] | -0.09 (-1.06 – 0.88) | 0.860 | -0.22 (-1.30 – 0.86) | 0.686 | -0.70 (-1.65 – 0.25) | 0.147 |
| WaterColor [White] | 0.23 (-0.59 – 1.06) | 0.573 | -0.71 (-1.67 – 0.24) | 0.138 | -0.32 (-1.15 – 0.52) | 0.455 |
| NetwD | 0.19 (-0.04 – 0.41) | 0.100 | 0.15 (-0.11 – 0.41) | 0.248 | 0.13 (-0.10 – 0.36) | 0.259 |
| Area | -0.09 (-0.40 – 0.22) | 0.554 | -0.15 (-0.47 – 0.16) | 0.340 | **-0.45 (-0.73 – -0.17)** | **0.002** |
| SoilDiv | 0.11(-0.16 – 0.38) | 0.412 | 0.18 (-0.11 – 0.47) | 0.216 | 0.15 (-0.10 – 0.41) | 0.239 |
| DistMouth | -0.82 (-1.87 – 0.23) | 0.123 | **-1.08 (-1.72 – -0.44)** | **0.001** | -0.51 (-1.07 – 0.05) | 0.076 |
| CoverDiv | 0.00 (-0.36 – 0.36) | 0.996 | 0.17 (-0.25 – 0.59) | 0.432 | -0.09 (-0.46 – 0.28) | 0.627 |
| PC1_Diff_CurrentLGM | -0.18 (-0.58 – 0.21) | 0.361 | -0.24 (-0.67 – 0.20) | 0.279 | -0.27 (-0.66 – 0.11) | 0.161 |
| PC2_Diff_CurrentLGM | 0.20 (-0.15 – 0.56) | 0.261 | **-0.41 (-0.79 – -0.04)** | **0.032** | 0.20 (-0.13 – 0.54) | 0.225 |
| PC3_Diff_CurrentLGM | -0.13 (-0.45 – 0.19) | 0.426 | -0.11 (-0.47 – 0.25) | 0.551 | -0.15 (-0.47 – 0.17) | 0.363 |
| PebasLake | 0.13 (-0.69 – 0.95) | 0.746 | -0.23 (-1.09 – 0.62) | 0.587 | 0.04 (-0.71 – 0.80) | 0.910 |
| Sea water at <1Mya | -0.11 (-0.49 – 0.26) | 0.553 | **0.51 (0.06 – 0.96)** | **0.026** | -0.17 (-0.56 – 0.22) | 0.384 |
| Sea water at ~5Mya | 0.20 (-0.17 – 0.56) | 0.283 | 0.39 (-0.03 – 0.81) | 0.069 | 0.01 (-0.36 – 0.38) | 0.950 |
| Waterfall | 0.00 (-0.26 – 0.26) | 0.996 | 0.14 (-0.15 – 0.43) | 0.348 | -0.01 (-0.26 – 0.25) | 0.962 |
| PC1_GlobEnv | 0.16 (-0.52 – 0.84) | 0.641 | -0.08 (-0.72 – 0.57) | 0.814 | 0.26 (-0.30 – 0.83) | 0.356 |
| PC2_GlobEnv | -0.32 (-0.78 – 0.14) | 0.166 | 0.34 (-0.12 – 0.80) | 0.142 | **-0.48 (-0.88 – -0.08)** | **0.020** |
| PC3_GlobEnv | 0.15 (-0.16 – 0.46) | 0.353 | 0.30 (-0.01 – 0.62) | 0.059 | 0.15 (-0.13 – 0.42) | 0.296 |
| PC4_GlobEnv | 0.25 (-0.03 – 0.53) | 0.085 | -0.02 (-0.30 – 0.26) | 0.884 | **0.25 (0.01 – 0.50)** | **0.043** |
| SamplingEffort | -0.06 (-0.33 – 0.22) | 0.684 | **-0.49 (-0.80 – -0.17)** | **0.003** | **-0.37 (-0.65 – -0.09)** | **0.010** |
| MEM [, 1] | 0.15 (-0.64 – 0.94) | 0.704 | | | | |
| MEM [, 2] | 0.16 (-0.38 – 0.70) | 0.556 | | | | |
| MEM [, 3] | -0.22 (-0.61 – 0.17) | 0.264 | | | | |
| *Moran's I (p value)* | -0.01 (p=0.33) | | -0.01 (p=0.38) | | -0.01 (p=0.37) | |
| $R^2$ / $R^2$ adjusted | 0.468 / 0.310 | | 0.740 / 0.676 | | 0.414 / 0.269 | |

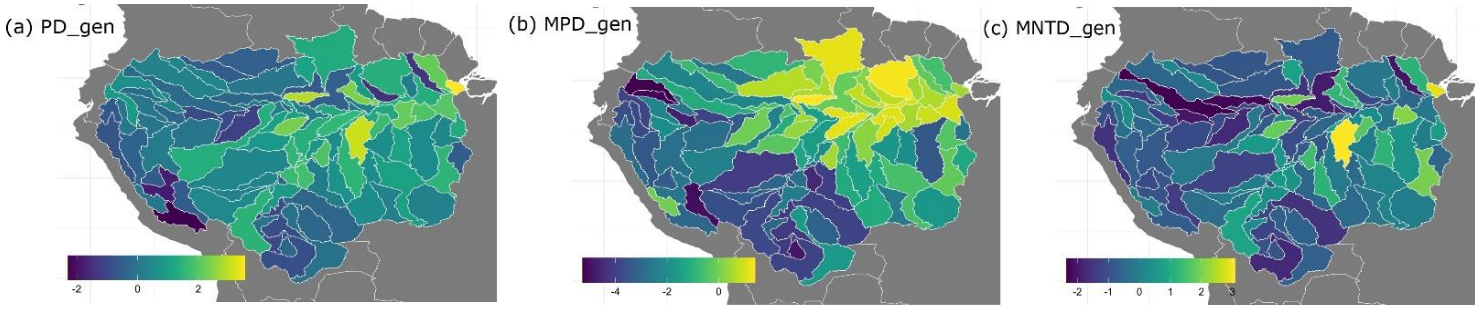


**Fig. S2.6.** Standardized effect size of phylogenetic diversity (ses.PD), mean phylogenetic distance (ses.MPD), and mean nearest taxon distance (ses.MNTD) calculated for 635 (genetic tree) native Amazonian freshwater fish species. Negative values of ses.MNTD and ses.MPD indicate phylogenetic clustered pattern, while positive ones, an overdispersed phylogenetic pattern.
